# Restoring the pattern of proteoglycan sulphation in perineuronal nets corrects age-related memory loss

**DOI:** 10.1101/2020.01.03.894188

**Authors:** Sujeong Yang, Sylvain Gigout, Angelo Molinaro, Yuko Naito-Matsui, Sam Hilton, Simona Foscarin, Bart Nieuwenhuis, Joost Verhaagen, Tommaso Pizzorusso, Lisa M. Saksida, Timothy M. Bussey, Hiroshi Kitagawa, Jessica C.F. Kwok, James W. Fawcett

## Abstract

Memory loss is a usual consequence of ageing and aged mice show progressive deficits in memory tasks. In aged brains, perineuronal nets (PNNs), which are implicated in plasticity and memory, become inhibitory due to decreased 6-sulphation of their glycan chains (C6S). Removal of PNNs or digestion of their glycosaminoglycans rescued age-related memory loss. Premature reduction of permissive C6S by transgenic deletion of chondroitin 6-sulfotransferase led to very early memory loss. However, restoring C6S levels in aged animals by AAV delivery or transgenic expression of 6-sulfotransferase restored memory. Low C6S levels caused loss of cortical long-term potentiation, which was restored by AAV-mediated 6-sulfotransferase delivery. The study shows that loss of C6S in the aged brain leads to declining memory and cognition. Age-related memory impairment was restored by C6S replacement or other interventions targeting perineuronal nets

## Introduction

Diminished ability to form and retrieve memories is a usual consequence of ageing, known as age-related memory impairment (ARMI). Current treatments to maintain and restore memory in aged individuals are currently inadequate. Various explanations for memory loss have been proposed, including axonal loss, epigenetic changes, decline in neurogenesis, synaptic loss, neuronal dysfunction, oxidative stress, loss of myelin, extracellular matrix and others (1–6).

Here, we demonstrate a role for perineuronal nets (PNNs), specialized structures in the CNS extracellular matrix (ECM) in age-related memory loss, and show that ECM manipulation can rescue memory. Previous work points to a key role for chondroitin sulphate proteoglycans (CSPGs) in memory, particularly the CSPGs associated with perineuronal nets (PNNs). During memory acquisition, new inhibitory inputs to GABAergic parvalbumin_+_ (PV^+^) neurons are formed (7–9). The somata and dendrites of most of these PV^+^ neurons are surrounded by PNNs, condensed extracellular matrix structures that surround synapses and are involved in the control of developmental and adult plasticity (10–14). In young animals digestion of inhibitory chondroitin sulphate glycosaminoglycan (CS-GAG) chains (the main effectors of PNNs) with chondroitinase ABC (ChABC), enables increased formation of inhibitory synapses on PV^+^ neurons (7, 15), and enhances memory acquisition and duration in rodents, as does transgenic attenuation of PNNs (16, 17). ChABC digests CS-GAGs, the sulphated sugar chains that are responsible for much of CSPG function. CS-GAG digestion and blocking have also restored memory in animals that model Alzheimer’s disease with pathology from mutant tau or amyloid-beta.. In these animals direct action on CS-GAGs with an antibody treatment to block inhibitory 4-sulphated CS-GAGs (C4S) or digestion with ChABC both restore memory to normal levels (18–20). Together these results demonstrate a role for PNNs in memory and memory restoration, and show that PNN CS-GAGs are the key component (21).

The binding properties and functions of GAGs are determined by their pattern of sulphation (22–24). This pattern changes with age; at birth CS-GAGs in the PNNs are 18% 6-sulphated (C6S), at the end of critical periods C6S declines to 4%, then in aged rats there is an almost complete loss of C6S (< 1%) (25, 26) leading to an increase in the ratio of C4S/C6S. C6S is permissive to axon growth and plasticity while C4S is inhibitory (27–29), so this change results in the PNN matrix becoming much more inhibitory in aged brains (25). For formation of new synapses onto PV^+^ neurons, processes must penetrate the PNNs; as these structures become more inhibitory with age, synaptogenesis that underlies formation of new memories may therefore be partially blocked. We hypothesised that a cause of age-related memory impairment is due to PNNs becoming more inhibitory with age caused by an increase in the ratio of C4S/C6S, leading in turn to memory loss associated with diminished inhibitory synapse formation onto PV^+^ interneurons. Our results support this hypothesis. We show that mice lose C6S with age, develop impaired memory. Memory can be restored by digesting CS-GAGs in the brain with ChABC. Transgenic animals with attenuation of PNNs show no age-related memory impairment.

Manipulation of the C6S levels in the brain leads to changes in memory, with lowered C6S leading to very early memory loss but enhanced C6S leads to prevention of ARMI and memory restoration. Restoration of C6S also restored LTP in the aged hippocampus and cortex. These memory events correlate with changes in the number of inhibitory synapses on PV^+^ interneurons. The study indicates that treatments targeting PNNs will be an effective means of restoring lost memory and preventing ARMI.

## Results

### Aged mice have memory deficits

We first established that aged 20 month-old (20M) mice have a progressive memory and cognitive deficit compared to young 6 month-old mice (6M) using three tests. These were a) spontaneous object recognition (SOR) (which depends on the perirhinal cortex (PRh), allowing focal application of treatments), b) spontaneous alternation (SA) (which tests hippocampal-dependent spatial memory), and c) marble burying (MB) (a test of exploratory response to novel environments, related to hippocampal function (30)).

From 20M C57BL/6 mice showed defective retention of SOR: normally young animals show memory of the novelty of objects at 6 hours (hr), forgetting by 24 hr (Fig.1A). In aged 20M animals SOR memory was barely detectable at 6 hr while 6M animals showed robust memory. However aged animals were able to form short-term memories similarly to young animals, detected by testing the memory within a minute of object exposure (Fig.1A). Participation in the tasks remained normal (Fig.S1A) indicating cognitive decline rather than a motor or attention deficit. SA was normal in 6M mice, but at 20M an abnormally high rate of same arm re-entry was seen (Fig.1B). In the MB test, mice behaved normally at 4 and 7M, but at 12 and 20M they buried abnormally few marbles (Fig.1C).

**Fig. 1.**
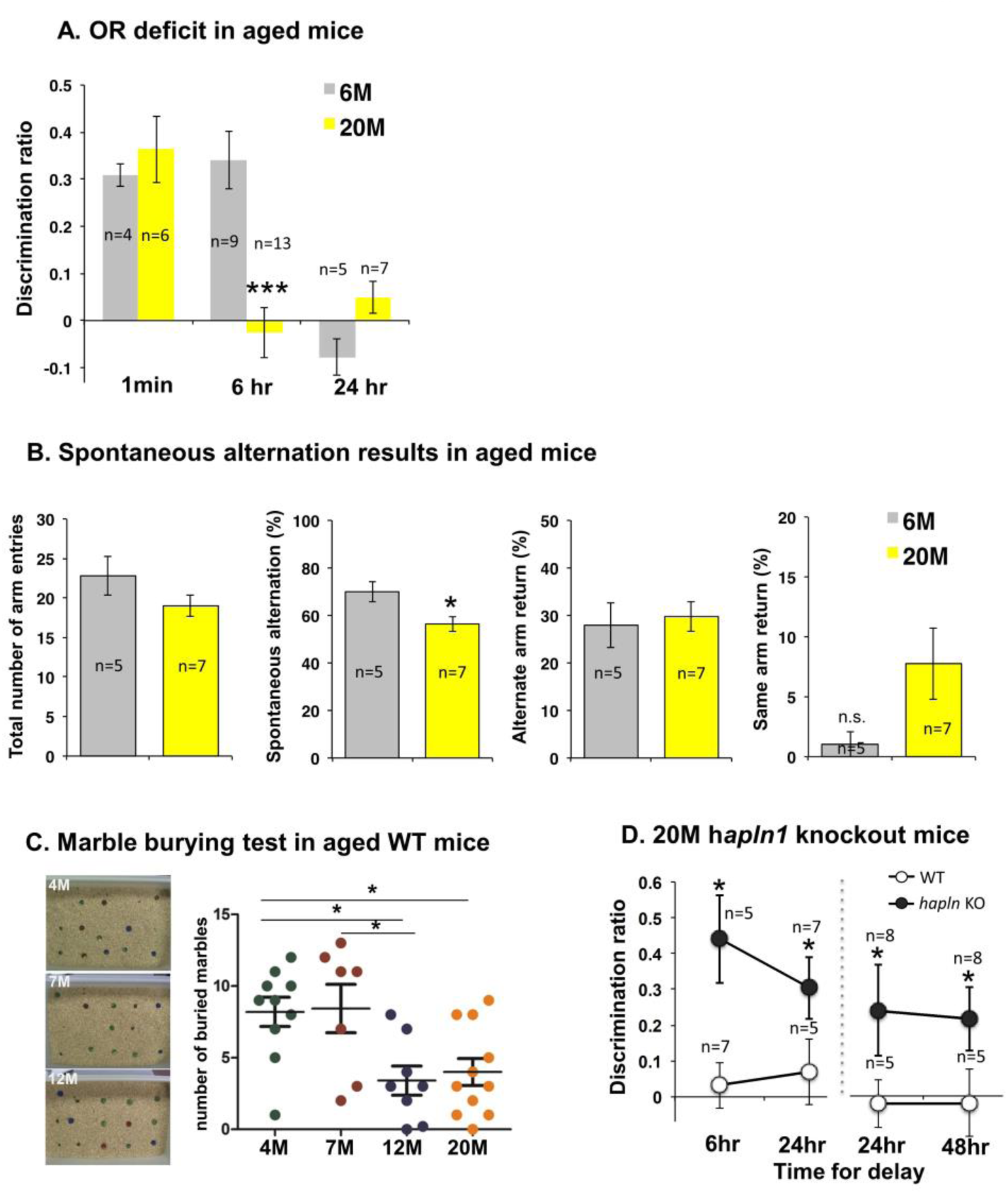
Memory loss in aged mice, rescued by perineuronal net attenuation. (A) OR memory deficit in aged mice. Unpaired two-tailed t-test, ***p=0.0002 (C) 6M vs 20M Low PV: **p=0.0012, High PV **p=0.0061, n=6/group, Data present as mean ± SEM. (B) Spontaneous alternation deficit in aged mice. Data is mean ± SEM. (C) Marble burying test in C57bl/6 at different ages, Left: representative images of marble burying test Right: 4M n=10, 4M n=6, 7M n=8, 20M n=11, One-way ANOVA **p=0.004, Tukey post hoc test, 4M vs 12M *p<0.05, 4M vs 20M *p<0.05, 7M vs 12M *p<0.05, Data represents as mean ± SEM. (D) SOR memory in aged *hapln1* KO mice (two separate experiments). Unpaired two-tailed t-test. 6hr delay: *p=0.0094. 24hr delay: *p=0.032, *=p=0.0226. 48hr delay: *p=0.0145 48hr delay: *p=0.0295.

### Transgenic attenuation of PNNs prevents age-related memory loss

To investigate if memory loss in ageing is related to PNNs, we assessed memory loss using SOR in transgenic animals with attenuated PNNs. One of the key PNN molecules is hyaluronan and proteoglycan link protein 1 (*hapln1)* which stabilizes the binding of CSPGs to hyaluronan and is necessary for the formation of stable condensed matrix structures like the PNNs. Transgenic deletion of *hapln1* leads to marked attenuation of PNNs, but brains contain normal quantities of CSPGs which are now diffusely spread (26, 31). In previous work we have shown that SOR memory is prolonged and various forms of plasticity are enhanced in non-aged *hapln1* knockout mice (17, 26). To investigate the role of PNNs in memory loss in ageing, we tested SOR in *hapln1* knockout animals at time points from 12M up to 20M. Even at 20M, these animals showed no loss of SOR memory, with memory retention extended to 48 hr, while littermates that expressed *hapln1* showed complete memory loss (Fig.1D). This result implicates PNNs in the loss of SOR memory in aged mice.

### Chondroitinase digestion of CSPGs restores memory

The effects of PNNs are dependent on their content of CSPGs (32). To investigate a link between CSPGs and memory loss in the aged brain, we used ChABC to digest CS-GAGs in the PRh which regulates SOR memory. ChABC was injected focally into the PRh, as in previous experiments that have assayed the effects of CSPGs on memory (17). An injection of ChABC into the PRh of 20M animals completely restored 6 hr memory retention, but did not lead to the abnormally extended memory seen after ChABC injections in young animals, where animals still recognise objects at 24 hr after exposure (Fig.2B). Following this restoration of memory, the memory declined again as the digested CSPGs were re-synthesised. By 6 weeks after treatment memory was back to baseline (timeline Fig.2A, Fig.2B). SA and MB (which do not depend on PRh) were not affected by this focal treatment and remained impaired. The results indicate that CS-GAGs in the PRh are responsible for the memory loss in the aged mice.

**Fig. 2.**
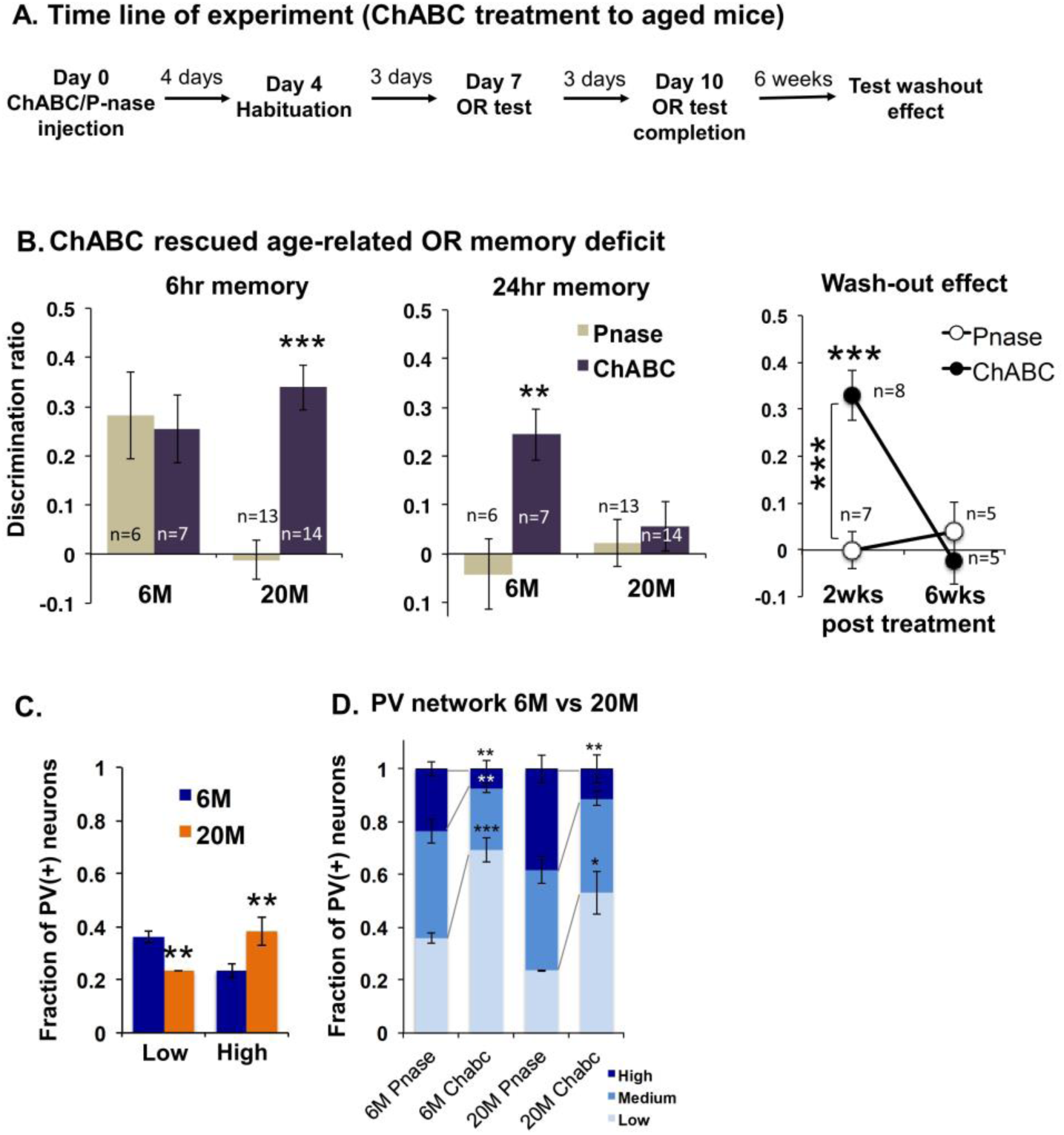
Chondroitinase ABC treatment restores memory and Parvalbumin levels. (A) Timeline. (B) ChABC injection rescued age-related OR memory deficit. 6hr memory, unpaired two-tailed t-test, *** p=0.0001. Wash-out of ChABC treatment effect of 6 hr memory in 20M mice. 2-Way ANOVA interaction treatment × time, ***p<0.001. Main effect of Time: ** p<0.01. Treatment: ** p<0.01. Bonferroni posthoc test, ChABC 2wks vs 6wks, ***p<0.001. 2wks Pnase vs ChABC, ***p=0.0002. (C) 6M vs 20M Low PV: **p=0.0012, High PV **p=0.0061, n=6/group, Data present as mean ± SEM. (D) PV network modification after ChABC treatment in 6M vs 20M old mice 6M Pnase vs ChABC n=6/group, Low: ***p<0.0001 Medium: **p=0.0027 High: **p=0.0028, 20M Pnase vs ChABC n=4/group, Low: *p=0.019 High: **p=0.0099

In young animals increased numbers of inhibitory synapses onto PV^+^ interneurons are seen during memory acquisition: this increased level of inhibition leads to a decrease in their PV and GAD67 levels, indicating decreased GABA production (7). ChABC-mediated CS-GAG digestion in young animals similarly increases inhibitory inputs and decreases PV expression (7, 33). We asked whether changes in the number of inhibitory synapse are seen in aged brain, and whether inhibitory synapse number is restored by ChABC digestion. We therefore measured the proportion of PV^+^ interneurons in untreated and ChABC-treated PRh. We counted PV^+^ neurons expressing high, medium and low levels of PV as in these previous studies. At 20M we saw an increase in the proportion of high-expressing PV^+^ cells and a decrease in low-expressing PV^+^ neurons in the aged cortex compared to 6M (Fig.2C). ChABC treatment at both ages increased low-expressing and decreased high-expressing PV^+^ neurons (Fig.2D, Fig.S1D). The overall number of neurons and PNNs in PRh was unchanged during ageing (Fig.S1B). These experiments show that SOR memory in aged animals can be restored by digestion of CS-GAGs in PRh. Together with the results from the link protein (*hapln 1*) knockouts, the conclusion is that CS-GAGs in PNNs are responsible for age-related memory impairment. Ageing is associated with increased PV expression in PV^+^ interneurons, levels being returned to those of young animals by ChABC treatment.

### Transgenic reduction of C6S leads to premature memory loss

We next investigated the changes in PNN CS-GAGs that are responsible for age-related memory loss. Our hypothesis was that the loss of C6S, causing an increased ratio of C4S/C6S in the aged brain, leads to CSPGs becoming increasingly inhibitory, leading in turn to defective memory in the aged brain. The existing data on changes in the sulphation levels of CS-GAGs with age was from rats (25), so we measured C4S and C6S in young and aged mice. The mouse results were very similar to rats: we found a reduction of C6S in aged 20M mice, leading to an increase in the ratio of C4S to C6S by 56% in PNN extracts (Fig.3A).

If loss of C6S is a causal factor in age-related memory loss, a decrease in C6S at any age would be expected to lead to memory impairment. To test whether premature C6S loss affects memory, SOR, SA and MB were tested in C6 sulfotransferase-1 (*chst3*) knockout animals, in which C6S is greatly diminished, C4S levels are normal and the ECM is abnormally inhibitory (29). At 8 weeks of age (8W), SOR memory formation and retention were normal, but by 11W there was a clear deficit, and by 13W there was no measurable memory retention even at 3 hrs after exposure to the objects (Fig. 3B). These animals also showed defective spatial memory as measured by SA. By 4M of age, the knockout animals showed increased frequency of immediate re-visiting the same arm of the maze (Fig. 3C, Fig. S2B). The animals also showed defective marble burying at 12M (Fig.3D). The number of PNNs was normal by WFA staining in these mice (Fig.S2C).

**Figure 3.**
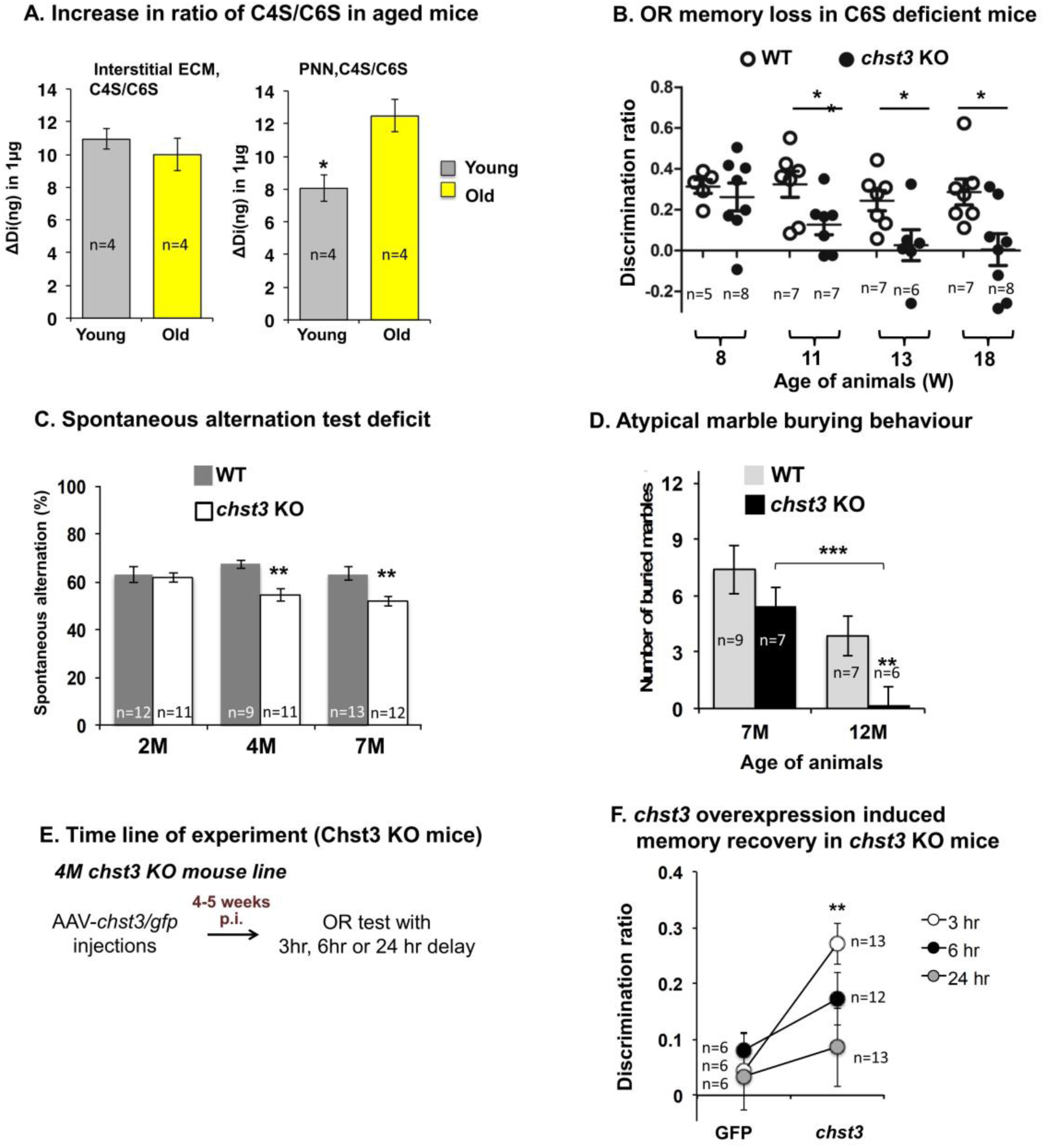
Aged mice lose C6S. Knockout of C6 sulfotransferase levels lead to premature memory loss. (A) PNNs in aged mice are depleted of C6S. C4S/C6S ratio in young and aged mice. Left general interstitial matrix, Right PNN fraction (B) OR memory loss (3 hr) in *chst3* knockout mice. Unpaired two-tailed t-test. 11 weeks: *p=0.0317. 13 weeks: *p=0.0306. 18 weeks: *p=0.0165. (C) Impaired spontaneous alternation performance in *chst3* knockout mice. Unpaired two-tailed *t*-test, 4M: **p=0.0011. 7M: **p=0.001. (D) Atypical marble burying behaviour in *chst3* KO mice on ageing. Unpaired two-tailed t-test. 12M: **p<0.01. In WT the aging impaired marble burying behaviour, 7M vs 12M ***p<0.001 (E) Timeline, (F) *chst3* gene delivery recovered memory in 4M *chst3* KO mice line. Unpaired two-tailed *t*-test, 3hr delay: **p<0.01.

In order to confirm that the loss of memory was due to changes in C6S levels and not due to an idiosyncrasy of the transgenic, we tested whether restoration of C6S could recover memory in *chst3* knockout mice. An AAV1 vector expressing mouse *chst3* under a PGK promoter was injected into the PRh at 4 months of age (Fig.3E). Injected brains were stained with the anti-C6S antibody CS56 (34), demonstrating an increased C6S level in the injected region associated with PNNs and other structures. In these AAV-*chst3-* injected knockout animals memory was restored to normal, demonstrating that the memory deficit of the knockout can be corrected by restoring the level of C6S in the PRh (Fig.3F). This result shows that rendering brain CS-GAGs abnormally inhibitory by decreasing the level of C6S leads prematurely to defective object recognition and spatial memory, and decreased marble burying behavior in adult animals.

### Restoration of C6S levels rescues age-related memory impairment

We next tested whether expression of *chst3* to reinstate the level of C6S would rescue memory in aged normal animals. Young and aged groups of C57BL/6 animals (5M and 19M) were tested for SOR, demonstrating a profound memory deficit in the aged animals as described above.

These elderly animals and young animals were injected with AAV1*-chst3* into PRh. 5 weeks later animals were tested for SOR memory at 6M and 20M of age (timeline Fig.4A). In the aged animals there was restoration of SOR memory at the 3 and 6 hr time points almost to levels normally observed in young animals while 3 and 6 hr memory was unaffected in young animals (Fig.4B). Young 6M animals injected with AAV1-*chst3* showed changes very similar to those caused by ChABC injection, with memory persisting for 24 hours (Fig.4B). There was no effect on SA or MB in these injected animals, as expected because only PRh was affected by the virus injections. C6S levels measured by CS56 immunostaining were increased in the injected region (Fig. S3C).

We next investigated the effects of a global increase in C6S. Transgenic animals constitutively overexpressing *chst3* have previously been shown to have enhanced plasticity as adults (35). In these animals the increased C6S is present throughout the brain, contrasting with the focal increase caused by injection of the AAV1-*chst3* virus. We asked whether these C6S overexpressing mice are protected from ARMI. At 20M these *chst3* overexpressing transgenic mice showed no memory deficit, possessing normal SOR 6hr memory while the littermate group showed a memory deficit similar to the aged animals described above (Fig. 4C). Because the *chst3* expression in these transgenic mice is general rather than focal, we also tested SA memory and MB behavior, and found that the age-related decline in these tests was also prevented (Fig. S3D,E).

**Fig. 4.**
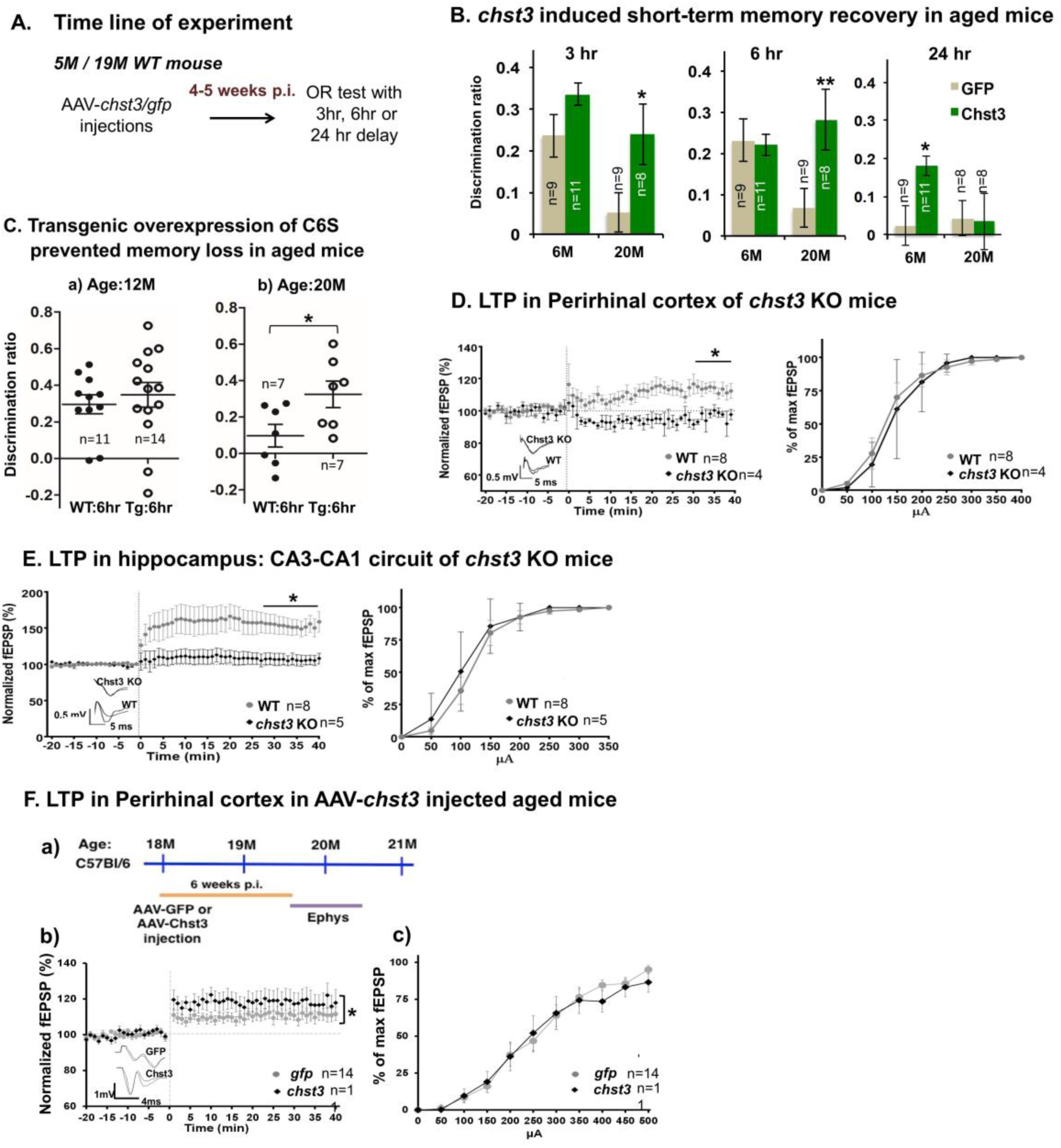
ARMI is prevented in animals overexpressing C6 sulfotransferase. Restoration of C6S restores LTP in brain slices. (A) Timeline (B) Age-mediated memory loss in C57bl/6 mice and SOR memory restoration by injection of AAV1-*Chst3* to PRh. 3hr memory retention: *p=0.023. 6 hr memory retention: **p=0.0071. (C) In *chst3* transgenic mice age-mediated memory loss was absent. Age: 20M, 6hr memory retention: WT vs Tg *p=0.0361. Data present as mean ± SEM. (D) LTP in perirhinal cortex slices in *chst3* KO mice. Left: LTP is significantly decreased (two-way RM ANOVA: *p=0.018). Left inset: Examples of fEPSP. Right: I/O curve of evoked fEPSP amplitude (two-way RM ANOVA, n.s. p=0.758;) (E) LTP is increased in CA3-CA1 in *chst3* KO mice (two-way RM ANOVA, *p=0.010). Left inset: Examples of fEPSP. Right: I/O curve (two-way RM ANOVA, n.s. p=0.457). Graphs are mean ± SEM. (F) LTP in perirhinal cortex in AAV-*chst3* injected aged mice. a) Time line. b) LTP in perirhinal cortex (two-way RM ANOVA: *p=0.036). Left inset: Examples of fEPSP. c) I/O curve (two-way RM ANOVA, p > 0.05).

The effects of ageing and manipulation of C6S on electrophysiological measures of plasticity were examined. Long-term potentiation (LTP) and input/output curves were observed in acute slices from the PRh and hippocampus. C*hst3* knockout animals, which have low C6S and show premature memory loss, demonstrated a complete loss of LTP in layer II/III following temporal cortex stimulation and a similar loss in the hippocampus (Fig.4D and E, left). The input/output relation was not changed (Fig.4D and E, right). In aged animals, LTP in the cortex was similarly diminished (Fig.4F). However in aged animals whose PRh was injected 5 weeks previously with AAV-*chst3*, LTP was restored almost to the level of 3-6 month old animals, but with no change in the input/output curve (Fig. 4F). These results are consistent with an overall higher level of inhibition in animals with low C6S levels, with previous evidence of PNN control of cortical excitability (36), and with previous evidence of increased inhibition in aged cortex (37, 38). These results strongly support the hypothesis that the level of C6S has a profound influence on memory, that the age-related decline in C6S is a factor in age-related cognitive loss and that manipulation of 6-sulfotransferase activity could be a successful way to restore memory to the aged.

### Manipulation of C6S increases inhibitory synapses on PV^+^ interneurons

Previous work has shown that during memory acquisition PV levels in PV^+^ interneurons are decreased, probably caused by an increase in the number of inhibitory inputs to PV^+^ cells (7). Is a similar mechanism driving age-related memory impairment? We show above that ageing influenced PV levels, with an increase in the number of PV^+^ interneurons that express high levels of PV (high expressers) and a decrease in those expressing low levels of PV (low expressers).

Also in our experiment above, treatment of PRh with ChABC (Fig. 2D) led to a decrease in PV (similar to result described by Donato *et al.* (7)), and enhanced SOR memory. We therefore asked whether during memory restoration by AAV-*chst3* injections there would be a restoration in the number of inhibitory synapses contacting PV^+^ interneurons. AAV-*chst3* was injected into PRh and after 5 weeks PV levels and synapse numbers were measured by immunostaining and PNNs were assayed by WFA lectin staining. In the AAV-*chst3-*injected PRh, the number of high PV-expressing neurons decreased, the number of low-expressing neurons increased, similar to the effect of ChABC treatment (Fig.5Aa). Overexpression of *chst3* additionally lowered WFA lectin staining of PNNs (Fig. 5Ab). We asked whether the number of inhibitory synapses on PV^+^ interneurons was changed. In aged animals the overall number of gephyrin+ synapses on PV cells was decreased compared to 6M animals, while treatment with AAV-*chst3* to restore C6S levels and memory led to restoration of the number of gephyrin synapses towards the level of young animals (Fig. 5B). Plotting PV levels *versus* number of gephyrin+ synapses for individual neurons revealed a close inverse correlation between the number of between the number of gephyrin+ synapses on PV^+^ cells and their level of PV expression (33). The slope of this correlation was greater at younger ages than in aged animals (Fig.5Bc). These results suggest a mechanism for age-related memory impairment. Aged PNNs lose C6S and become more inhibitory. This leads to a decrease in inhibitory inputs to PV^+^ neurons, causing increased PV expression and increased cortical inhibition. The increased GABAergic inhibition impairs memory acquisition. Removing PNNs or restoring C6S levels enables increased inhibitory synapses on PV^+^ interneurons, facilitating restoration of normal memory.

**Fig. 5.**
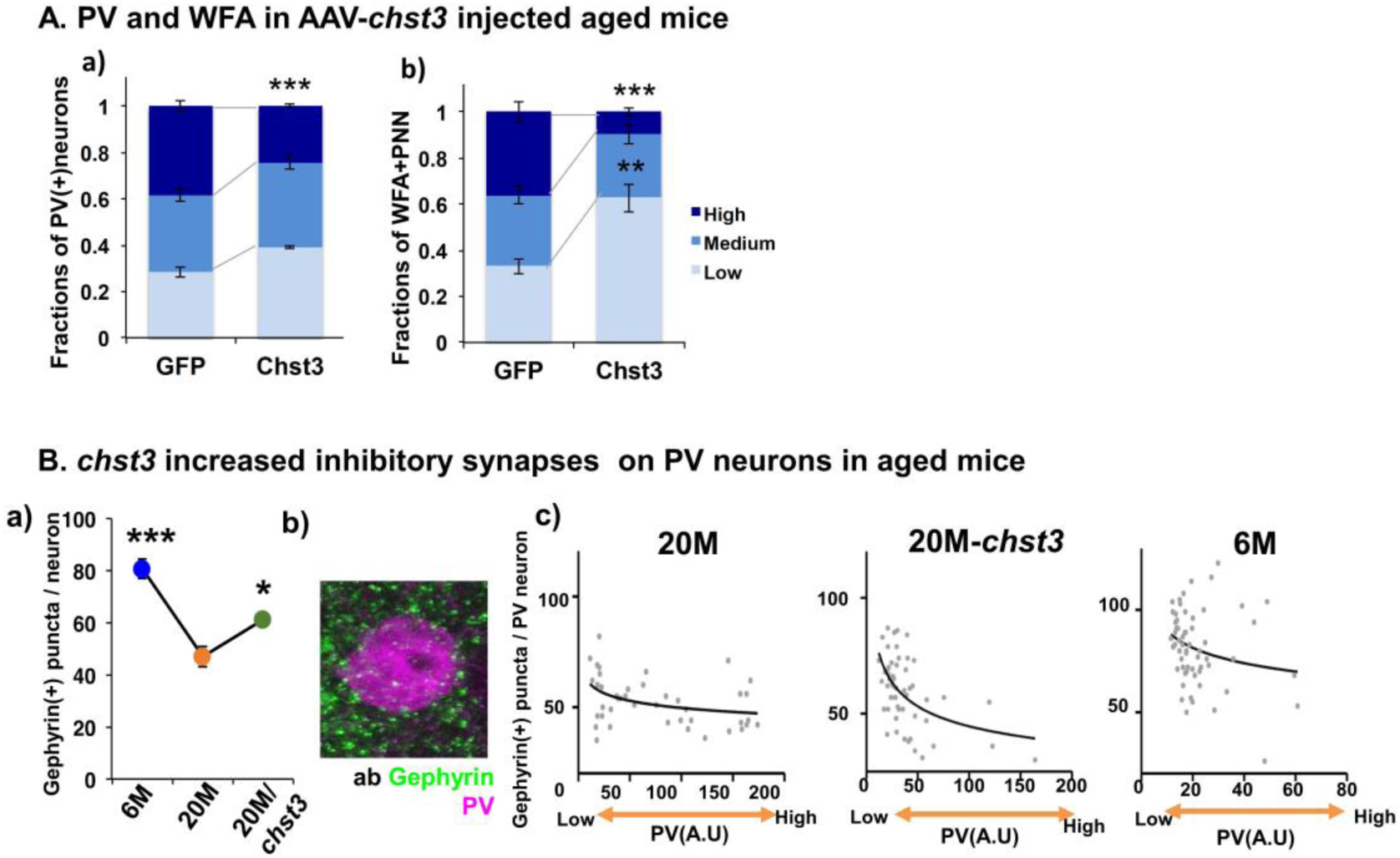
Relation between C6S and long-term potentiation (LTP) (A) PV and WFA network in AAV-*chst3* injected aged mice a) PV network, High: ***p=0.0001 Low: n.s. p=0.079. b) WFA network, High: ***p<0.0001 Low: **p=0.0016. (Bab) *chst3* increased the inhibitory synapses on PV neurons in aged mice. a) n=6/group. 6M vs 20M ***p=0.0002. 20M vs 20M-*chst3*, * p=0.01. Unpaired two-tailed t-test. b) Staining profiles of gephyrin (+) synaptic puncta on PV cells. (Bc) Correlation profiles between numbers of gephyrin synaptic puncta and PV intensity. 20M: Correlation *p=0.0186, Spearman r=-0.3576, 20M-Chst3: Correlation ***p=0.0003, Spearman r= -0.4717, 6M: Correlation *p=0.0364, Spearman r=-0.2686

## Discussion

In the current study we asked whether PNNs and their constituent CSPGs, particularly the sulphation composition of the CS-GAGs, are involved in ARMI, and whether their manipulation can restore memory and prevent ARMI. We used three behavioural models: two models of memory (SOR and SA) and one of species-specific exploratory behaviour (MB) all showed an age-related deficit. PNN function depends on the CS-GAGs of CSPGs. Focal injection of the CS-GAG-digesting enzyme ChABC to the PRh of aged mice restored SOR, but did not affect SA and MB which rely on different brain areas which were not digested. However, an effect of ChABC does not in itself implicate PNNs, because ChABC digests CSPGs both in PNNs and in the diffuse CNS extracellular matrix. To demonstrate a specific role of PNNs in ARMI, transgenic animals with a deletion of *hapln1,* (a link protein that stabilizes the interaction of CSPGs and hyaluronan in PNNs) was used. These animals have very attenuated PNNs, but normal overall levels of CSPGs (26). When these PNN-attenuated animals were allowed to age they showed no age-related memory loss. In addition, we found that the down-regulation of C6S is responsible for ARMI. Transgenic mice with *chst3* over-expression or viral expression of *chst3* lead to restoration of C6S level to juvenile level and rescue ARMI. These results imply that the CS-GAGs of CSPGs in PNNs are involved in ARMI, and that manipulating these elements can reverse or prevent ARMI.

Age-related memory impairment (ARMI) affects the majority of elderly people. It is distinct from Alzheimer’s disease, which is more severe and more rapidly progressive and has a different pathology. CNS plasticity, including memory is strongly influenced by the extracellular matrix, particularly by the perineuronal nets (PNNs), condensed cartilage-like structures that mostly surround PV^+^ inhibitory interneurons in the brain. PNNs are involved in the closure of critical periods for plasticity (10, 14, 21, 39). Attenuation or digestion of PNNs restores many forms of plasticity. CSPGs and PNNs have been implicated in several forms of memory. Digestion or transgenic attenuation of PNNs in normal animals prolongs object recognition memory, enables unlearning of stressful memories and addiction and enables social memory (17, 40-42). CSPG and PNN manipulation can also restore memory lost due to pathology in tauopathy and Abeta models of Alzheimer’s disease (20, 43, 44).

The inhibitory function of CSPGs relies mainly on their sulphated CS-GAG chains, with different forms of sulphation having different properties. Overall, 4-sulphated CS-GAGs (C4S) are inhibitory to regeneration and plasticity while C6S is inhibitory (27, 29, 35). The sulphation pattern in the CNS changes at key times, with the ratio of C4S/C6S increasing after embryogenesis and further at the end of critical periods for plasticity (26, 28). PNN CS-GAG sulphation finally changes with ageing with the loss of C6S, rendering the PNN GAGs more inhibitory (25, 45). From the evidence for PNN involvement in ARMI and the ageing-related changes in PNN CS-GAGs came the hypothesis that the increasing ratio of C4S/C6S with ageing, making PNNs more inhibitory, contributes to the development of ARMI. The hypothesis predicts that increasing or decreasing C6S through manipulation of the main CS-6-sulfotransferase *chst3* would have effects on memory and ARMI. First, animals with transgenic deletion of *chst3* were tested for the timing of memory loss. These animals showed early impairment in SOR, SA and MB, and SOR was restored by injecting an AAV expressing *chst3* into the PRh to restore the level of C6S. This implies that permissive CS-GAGs are needed throughout life in order to allow for age-related effects that threaten normal memory. The hypothesis predicts that restoration or preservation of C6S levels should enable better memory in aged animals. We tested this prediction by injecting AAV-*chst3* into the PRh, and testing transgenic animals that overexpress *chst3. Chst3* injection into PRh leads to a focal increase in C6S, and this restored SOR (which relies on the PRh) in aged animals. A transgenic animal with a global overexpression of *chst3* was also tested, and this showed a general halt to ARMI, with normal SOR, SA and MB in aged animals. Together these results show that the loss of C6S in PNNs is involved in ARMI, and restoration of C6S levels can halt ARMI and restore memory in aged animals.

What might be the mechanism by with the sulphation of CS-GAGs in PNNs affects memory? Previous work has shown that memory acquisition involves an increase in the number of inhibitory synapses on PNN-bearing PV^+^ interneurons, leading to decreased GABA production. Moreover ChABC digestion both enabled memory and increased synapses on PV^+^ interneurons (7, 33, 46). The implication for ARMI is that the inhibitory PNNs that appear with ageing might block new synapse formation of PV^+^ interneurons, so inhibiting memory acquisition. The results support this hypothesis, because the number of inhibitory synapses on cortical PV^+^ neurons is decreased in aged brains and the PV levels in PV^+^ interneurons are increased. The interventions that restore memory, ChABC and increased *chst3,* both restore inhibitory synapse number on PV^+^ neurons. The PV levels in PV^+^ interneurons indicate the level of GABA-producing GAD-67 (33), and therefore suggest an age-related increase in overall GABA-mediated cortical inhibition.

Ageing has been associated with a deficit in short-term plasticity in the CNS, as measured by a decline or loss of LTP. Physiology of acute cortical slices showed defective LTP in our aged brains, and LTP was restored in animals that had received AAV-*chst3* injections. How might changes in overall levels of inhibition mediated by PV^+^ GABAergic interneurons and influenced by the properties of PNNs affect memory acquisition? It is possible that each PV^+^ neuron controls a particular cortical circuit which can be activated to store a particular memory. However PV^+^ interneurons generally change activity collectively, and for patterns of individual PV^+^ cells to define a memory seems unlikely. A more probable idea comes from investigating the number and distribution of cells that form a memory engram. Here the strength and sustainability of a memory may depend on the number of distributed neurons involved, which in turn would be influenced by overall excitability (47, 48).

Memory is complex, and several age-related processes are probable participants in ARMI (2–5). It is unlikely that changes in PNN sulphation are the only cause or ARMI. However an attraction of this mechanism from the perspective of future therapeutics is that PNNs are rich in potential therapeutic targets, including the sulfotransferase enzymes, the production of the hyaluronan backbone of the PNNs (49), the maintenance of PNNs by the diffusible transcription factor OTX2 (50). ChABC is not a useful therapeutic for ARMI, because its range and duration of action are limited, but an anti-C4S antibody has proven successful at memory restoration. Overall the results of this study demonstrate a mechanism for the loss of memory in the aged brain, and indicate that treatments targeting PNNs have the potential to ameliorate memory deficits associated with ageing.

## Materials and Methods

### Mice

The normal wild type (WT) C57BL/6J (Charles river, UK) mice were used for ChABC treatment study and 6M of age represents for young adults and 20M for aged group. In order to test between C6S and memory, two transgenic mouse models were used; c6st-1 (encoded by chst3 gene) knockout mice (21) and chst3-1 overexpressing Tg mice (22). Age-matched littermates were used as control mice. Animals had unrestricted access to food and water, and were maintained on a 12 hr light/dark cycle (lights off at 7:00 P.M.). All experiments were carried out in accordance with the UK Home Office Regulations for the Care and Use of Laboratory Animals and the UK Animals (Scientific Procedures) Act 1986.

### Generation of adeno-associated viral vectors

The plasmid encoding AAV-PGK-chst3 was made by amplifying the mouse chst3 sequence from plasmid MR207541 (OriGene) via the primers 5’ GGAATTCATAGGGCGGCCGGGAA 3’ and 5’ AGCGCTGGCCGGCCGTTTAAAC 3’ and was cloned into plasmid AAV-PGK-Cre (Addgene plasmid # 24593) between the AfeI (NEB, R0652) and EcoRI (NEB, R0101) sites to substitute the Cre recombinase gene. The eGFP sequence of AAV-CMV-eGFP (Addgene plasmid # 67634) was amplified by using the primers 5’ GGAATTCATGGTGAGCAAGGGCGAG 3’ and 5’ AGCGCTTTACTTGTACAGCTCGTCCATG 3’, which was next cloned into the digested AAV-PGK-backbone. These virus vectors were turned into a recombinant adeno-associated viral vector with serotype 1 as described in previously published protocol (J. Verhaagen *et al.*, 2018). For the present study, the following vectors were produced: AAV1-PGK-*chst3* 1.44×10^12^ gc/ml; AAV1-PGK-GFP 1.42×10^12^ gc/ml; AAV1-SYN-GFP 8.99×10^12^ gc/ml

J. Verhaagen *et al.*, Small Scale Production of Recombinant Adeno-Associated Viral Vectors for Gene Delivery to the Nervous System. Methods Mol. Biol. 1715, 3–17 (2018).

### Animal surgeries

Animal surgeries were performed under isoflurane anesthesia. ChABC (50 U/ml in PBS, Seikagaku), or AAV vectors (AAV1-PGK-Chst3 or AAV1-PGK-GFPor AAV1-SYN-GFP) was stereotaxically injected to six different sites in the PRh (1×10^8^ particles in total, 3 per hemisphere, 0.5 μl with a speed of 0.2 μl/min). Injections were made with a 10 μl Hamilton syringe and a 33 gauge needle at the following sites (in mm from bregma and the surface of skull): 1. anteriorposterior (AP): -1.8; lateral (L): ± 4.6; ventral (V): -4.4. 2. AP: - 2.8; L: ± 4.8; V: -4.3. 3. AP: -3.8; L: ± 4.8; V: -3.8. The needle remained in place at the injection site for 3 min before being slowly withdrawn over 2 min. AAV1-SYN-GFP was injected as a control viral vector for chst3 gene delivery to chst3 KO mice.

### Spontaneous Object Recognition task

The spontaneous object recognition (SOR) task was performed as previously described for mice (Yang et al, 2015). Briefly, all mice were habituated in three consecutive daily sessions in the empty Y-maze apparatus for 5 min after recovery from surgery. Each test session consisted of a sample phase and a choice phase. In the sample phase, the animal was placed in the start arm and left to explore two identical objects, which were placed on the end of two arms for 5 min. The choice phase followed after a delay of either 1 min, 3 hr, 6 hr as a short-term memory paradigm or 24 hr and 48 hr as a long-term memory paradigm, which the animal spent in the home cage. The choice phase was procedurally identical to the sample phase, except that one arm contained a novel object, whereas the other arm contained a copy of the repeated object. When the animals were assessed, a different object pair was used for each session for a given animal, and the order of exposure to object pairs were counterbalanced within and across groups. All the sessions were recorded by a technician (SH) who is blind to the genotype. The object exploration time was assessed from video recordings of the sample and choice phase by scorer (SY) blind to the genotype of the mouse. The direct nasal or head contacts only were regarded as an exploratory behaviour. A discrimination ratio was calculated by dividing the difference in exploration of the novel and familiar objects by the total object exploration time. Therefore, the discrimination ratio varies from 0 (equal exploration for novel and familiar objects) to 1 (exploration of the novel object only). The test sessions were separated by a minimum of 48 hr. One-way analysis of variance (ANOVA) followed by Tukey post hoc test was used for comparisons of multiple groups with a significance level of p < 0.05, using GraphPad Prism version 5.0. An unpaired two-tailed t-test was used for two-group comparisons.

The minimum sample size per animal experiment required to have meaningful results was 14 animals, which was calculated based on raw data of chABC or control treated 20M old mice using following parameters; type 1 error rate (a):0.05 and power: 0.8, mu1 -0.002, mu 2 0.3301, sigma 0.214 (http://www.stat.ubc.ca/~rollin/stats/ssize/n2.html).

For some experiments two cohorts of mice were recruited and the results were analysed at the end of experiments.

### Spontaneous alternation test

The Y maze was made by three white, opaque Perspex plastic arms (8 cm width, 20 cm length, 35 cm height) at a 120° angle from each other. There were no visual cues inside the maze. Each animal was allowed to freely navigate all three arms for 5 min after placing it at the centre of multiple arms via a tube. The number of arm entries and the number of trials were recorded in order to calculate the percentage of alternation. An entry occurs when all four limbs are within the arm. The inside of Y maze was cleaned with 50% ethanol between trials and allowed to dry.

### Marble burying test

Standard polycarbonate rat cages (26 cm × 48 cm × 20 cm) with fitted filter-top covers was used as a testing chamber and fresh, unscented mouse bedding material to each cage to a depth of 5 cm. Glass toy marbles (assorted styles and colors, 15 mm diameter, 5.2 g in weight) were placed gently on the surface of the bedding in 3 rows of 5 marbles. Each animal was carefully placed into a corner of the cage containing marbles, as far from marbles as possible, and the lid was placed on the cage and remained for 30 min. Marble as buried if two-thirds of its surface area is covered by bedding was counted by a scorer blind to the genotype of the mouse.

### Immunohistochemistry

#### Diaminobenzidine (DAB) staining

Sample preparations and the general procedures of immunostaining have been described previously (Yang *et al.*, 2015). In general, 30 µm free floating sections were incubated with 10 % methanol, 3 % H_2_O_2_ in PBS for 20 min at room temperature for quenching endogenous peroxidase activity. Sections were rinsed three times in 0.2 % Triton-X in PBS (PBS-T) and were subsequently blocked with 5 % normal goat serum (NGS) or normal horse serum (NHS) in PBS-T for 1 hr at room temperature. The primary antibodies (PV; 1:1000, Swant; Biotin-WFA; 1:100, Sigma-Aldrich) were incubated over night at 4 °C following rinsing tissues for 5 min three times in PBS-T (0.2 % Triton-X in PBS). Following 3 washes in PBS-T they were incubated for 2 hr at RT with the appropriate biotinylated secondary antibody (Vector Laboratories) diluted 1:500 in PBS-T. Subsequently, sections were incubated with an avidin-biotin system (Vectastain; Vector Laboratories) for 1 hr at room temperature and washed with PBS-T. The immunostaining was visualized with DAB with 3 % H_2_O_2_ (DAB kit; Vector Laboratories) for 1-5 min at room temperature. The sections were mounted on gelatin-coated slides and air-dried. Following dehydrating tissue sections in ascending concentration of alcohols, they are cleared in xylene and coverslipped with DPX. The tissue sections were examined using a light microscope and photographed using a digital camera (Leica DM6000 Microsystems).

#### Fluorescent staining/Analysis

Sections were blocked with 5 % normal goat serum (NGS) or normal horse serum (NHS) in PBS-T for 1 hr at RT. The primary antibodies (PV; 1:1000, Swant CS56; 1:100, Sigma-Aldrich; Biotin-WFA; 1:100, Sigma-Aldrich, Gephyrin; 1:200, Synaptic system) were incubated overnight at 4 °C. Following 3 washes in PBS they were incubated for 2 hr at RT with the appropriate secondary antibody conjugated with Alexa fluor 647, Alexa fluor 488 or Alexa fluor 568 or Streptavidin-Alexa fluor 647 (Molecular Probes, Invitrogen) diluted 1:500 in PBS-T. incubated with secondary antibodies for 2 hours. Sections were rinsed and mounted on 1% gelatin coated slides with FluorSave™ Reagent (Merck Millipore, Germany).

For synaptic puncta quantification images were captured using a Leica SPE confocal microscope using x63 objectives with a 1024 × 1024 image resolution (n=6 per group). At least 3 z-stack images (total 5 µm) were taken per section with at least 3 sections analyzed per animal (approx. 360µm apart). Images contained at least 5 PV positive neurons. At least 50 PV^+^ neurons per animal were analysed for gephyrin (+) puncta quantification. Synaptic puncta analysis was performed with an automated custom script using an imageJ 1.29 plugin (available from) (Ippolito and Eroglu, 2010).

Ippolito DM, Eroglu C. Quantifying Synapses: an Immunocytochemistry-based Assay to Quantify SynapseNumber. J. Vis. Exp JoVE. 45, 2270 (2010).

#### GAG extraction

Biochemical analyses were performed on 5 animals at each age. Sequential extraction of GAGs from rat brains was performed according to the protocol from (Kwok et al. 2015). Briefly, fresh frozen brains were first homogenized sequentially in 4 buffers: buffer 1 [B1: 50 mM Tris-buffered saline (TBS), 2 mM ethylenediamine tetraacetic acid (EDTA)], buffer 2 (B2: 0.5% Triton X-100 in B1), buffer 3 (B3: 1 M sodium chloride in B2) and buffer 4 (B4: 6 M urea in B2). All buffers were supplemented with protease inhibitors. The homogenate was centrifuged at 23,000 xg for 20 min at 4°C after each homogenization. Supernatant was collected and the pellet was homogenized in the next buffer. Supernatants from each buffer were dialyzed in 25 mM Tris-HCl and 5 mM EDTA pH 8.0 overnight at 4°C and subsequently digested with 200 mg/ml pronase (Roche). Peptides were precipitated with 5% trichloroacetic acid (TCA) on ice, centrifuged and the supernatants were collected. Supernatants were then washed four times in 1:1 diethyl ether which was then aspirated. The samples, which contained the glycosaminoglycans (GAGs) were neutralized to pH 7.0 and precipitated with sodium acetate. GAGs were recovered from the pellets after centrifugation, air dried and resuspended in deionized water.

Recovered GAGs were quantified using cetylpyridinium chloride (CPC) turbidimetry assay. Chondroitin sulphate-A (Sigma Aldrich) was used to set up the standard curves.

#### Fluorophore-assisted carbohydrate electrophoresis (FACE)

Recovered GAGs were digested into disaccharides with 0.1 U chABC (Sigma), then precipitated in EtOH for 16 h at 4°C. For C6S quantification, the samples were further treated further with 50 mU of chondro-6-sulfatase (Seikagaku). Samples were centrifuged, supernatants were collected and dried in a SpeedVac. The dried pellets were derivatized with 2-aminoacridone (AMAC) in sodium cyanoborohydride containing buffer. 0.5 μg of AMAC-conjugated disaccharide samples and standard disaccharides (hyaluronic acid – HA, non-sulfated chondroitin - C0S, C4S, C6S; Seikagaku) were separated on a 30% polyacrylamide gel in Tris Glycine buffer for 30-40 min.

For C4S, gels were imaged in a UV chamber (UVitec). Bands size and intensity were quantified using ImageJTM software. The intensity of the sample was normalized against the intensity of non-sulfated CS (C0S) and the quantity of disaccharides was calculated using the standard curve of HA electrophoresed in the same gel.

#### Electrophysiology

Animals were anesthetized with an overdose of isoflurane, humanely killed by cervical dislocation and decapitated. The brain was rapidly removed and placed in ice-cold cutting solution bubbled with 99% O2 containing the following (in mM): 126 NaCl, 2.5 KCl, 1 CaCl2·H2O, 2 MgCl2, 1.25 NaH2Po4·H2O, 10 NaHCO3, 5 D-glucose, 0.4 ascorbic acid, 3 myoinositol, 3 pyruvate, and 15 HEPES, adjusted to pH 7.35. For the cutting and the recording of the perirhinal cortex slices from the 20M aged animal, we used a modified ACSF continuously oxygenated (carbogen: 95% O2-5% CO2) and containing the following (in mM): 124 NaCl, 3 KCl, 1.5 CaCl2·2H2O, 1.5 MgCl2 6H2O, 1.25 NaH2Po4·H2O, 26 NaHCO3, 10 D-glucose, 0.01 Glycine, 1 L-ascorbic acid, and 2 Na Pyruvate, adjusted to pH 7.35. For perirhinal cortex a midsagittal section was made and the rostral part of one hemisphere was cut at 45° to the dorsoventral axis (Cho *et al.*, 2000). The cerebellum was removed from the brain with a further caudal coronal cut. The hemisphere was glued by its rostral end to a Vibratome stage (VT 1000S; Leica). Slices (380 µm) of perirhinal cortex were taken in the region -2.5mm to -4mm rostral from bregma. For hippocampus the whole brain was glued by its caudal end and coronal sections were taken in the region of dorsal hippocampus. Slices were stored submerged in bubbled, artificial ACSF (20-25°C, same composition as cutting solution, except 2 mM CaCl2, 1 mM MgCl2 and without: ascorbic acid, myoinositol, pyruvate) for 2 hr before the onset of recordings. A single slice was placed in an interface recording chamber superfused by artificial CSF (30°C, flow rate 2 ml/min). Evoked field EPSPs (fEPSP) for perirhinal cortex were recorded from layers II/III from directly below the rhinal sulcus (area 35). A stimulation electrode was placed in layer II/III on the entorhinal side (0.5 mm, area 36) of the recording electrode (Cho *et al.*, 2000). fEPSP for hippocampus were recorder recorded placing the stimulation electrode in the stratum radiatum of the CA3 field and the recording electrode in the same layer of the CA1 field. Stimuli (0.1 ms duration) were delivered to the stimulation electrode at 0.1 Hz. Input/output curves were produced with stimulation intensities from 50 to 450 500 ?A in steps of 50 ?A. For monitoring baseline synaptic transmission before LTP induction, fEPSPs were reduced to 40-50 % of the maximum amplitude and recorded for at least 20 min or until responses were stable (< 20 % amplitude change over 30 min). For LTP induction, the following protocol was used: 4 bursts at interval of 15 s, each composed by 10 trains at interval of 0.2 s, each composed by 4 pulses at interval of 10 ms and of 0.2 ms of duration with 3 V of amplitude. Subsequently, fEPSPs elicited by stimulations at interval of 20 s were recorded for further 40 min. Field potentials were amplified with a CyberAmp 320 (Molecular Devices) or an Axoclamp-2B (Axon Instrument), and recorded and analyzed with pClamp (Molecular Devices) or a custom-made software written in LabView (National Instruments). For offline LTP analysis, fEPSPs were averaged across 1 min and the peak amplitude of the mean fEPSP was expressed relative to the preconditioning baseline. The significance of group means was established using repeated-measures (RM) ANOVA.

## Abbreviations

AAV: Adeno-associated virus
ARMI: Age-related memory impairment
C4S: chondroitin 4 sulphate
C6S: chondroitin 6 sulphate
ChABC: Chondroitinase ABC
CSAPG: Chondroitin sulphate proteoglycan
CS-GAG: chondroitin sulphate glycosaminoglycan
GAG: Glycosoaminoglycan
LTP: Long term potentiation
MB: Marble burying
PNNs: Perineuronal nets
PRh: Perirhinal cortex
PV: Parvalbumin
SA: Spontaneous alternation
SOR: Sponteneous object recognition
WFA: Wisteria floribunda agglutinin

## Author Contributions

Experimental work was performed by SY, SG, AM, YN-M, SH, SF, BN, JCFK. Advice and supervision was from JV, TP, LMS, TMB, HK, JWF. Writing of the manuscript involved all authors. Grant funding was to SY, JV, LMS, TMV, HK, JCFK, JWF.

## Acknowledgements

Funding for this study was a fellowship to SY from Alzheimer’s Research UK (ARUK-RF2016A-1), grants from European Research Council ECMneuro 294502, Medical Research Council G1000864, MR/R004463, MR/S011110/1, CiC, Alzheimer’s Research UK Yorkshire network, Czech Science Foundation GACR 19-10365S Center of Reconstructive Neuroscience (CZ.02.1.01/0.0./0.0/15_003/0000419)

---

**Supplementary Fig.1.**
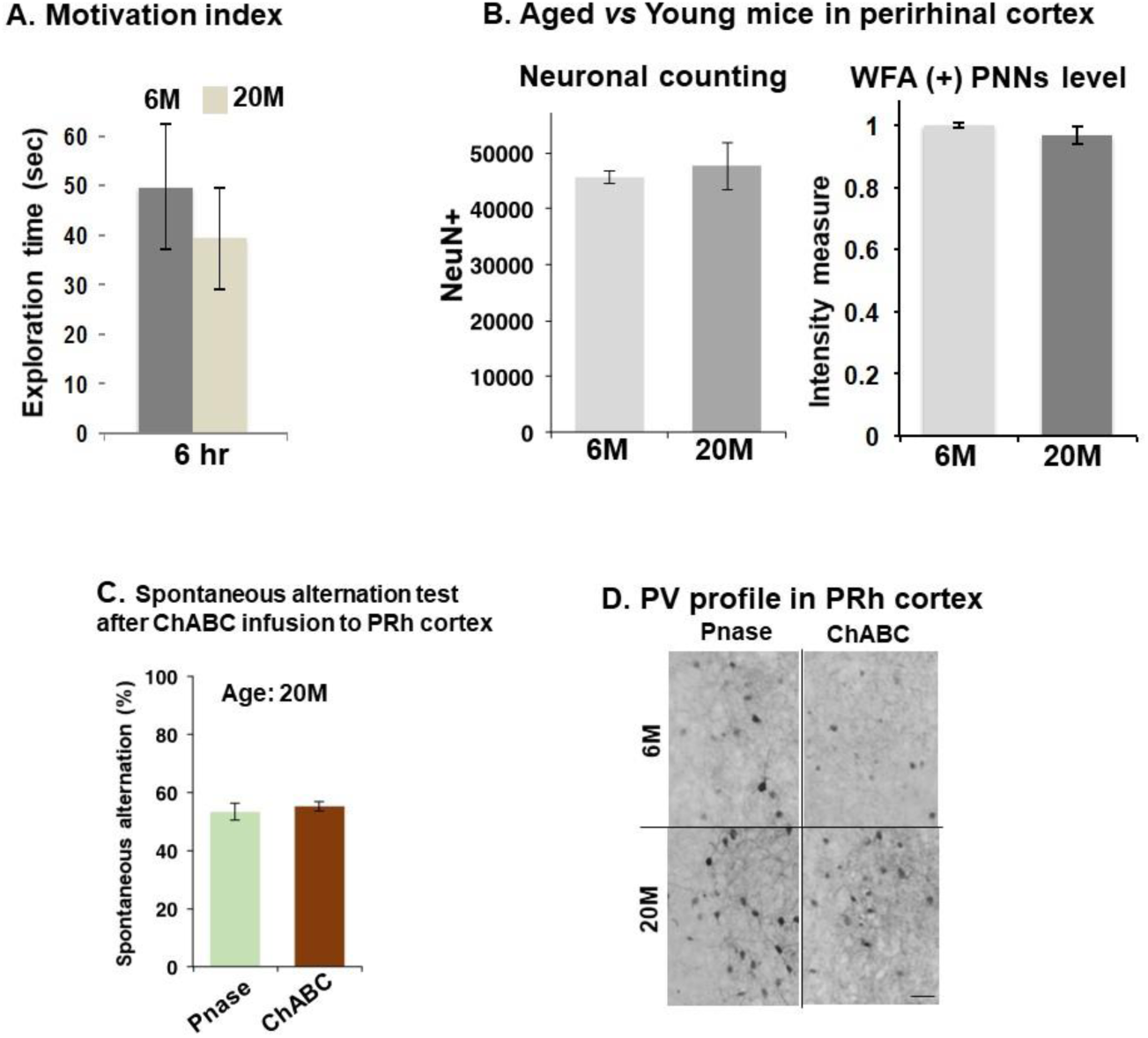
(A) Motivation index indicated by exploration time during sample phase of SOR test. No significant alteration in motivation was displayed between young and aged mice groups. 6M n=4, 20M n=7 (B) Neuronal counting (Left) and WFA(+) PNNs level (Right) in the perirhinal cortex of 6M or 20M old C57bl/6 mice. NeuN(+): n=6/group, WFA(+): n=4/group. (C) Marble burying test in C57bl/6 at different ages, Left: representative images of marble burying test Right: 4M n=10, 4M n=6, 7M n=8, 20M n=11, One-way ANOVA **p=0.004, Tukey post hoc test, 4M vs 12M *p<0.05, 4M vs 20M *p<0.05, 7M vs 12M *p<0.05, Data represents as mean ± SEM. (D) ChABC treatment to aged mice, 19M-old C57bl/6 mice were used. After 4 days of treatment, habituation of SOR test was performed and OR memory was measured at 7 days post injection. (E) ChABC or Pnase injections to perirhinal cortex. It is a schematic illustration of sagittal section of murine brain. a: bregma -1.82mm b: bregma -2.8mm c: bregma -3.8mm (F) Spontaneous alternation performance after ChABC treatment to perirhinal cortex (PRh) in aged mice. (G) Immunohistochemistry of PV in PRh cortex after ChABC or Pnase injection. Scale bar: 50 µm

**Supplementary Fig.2.**
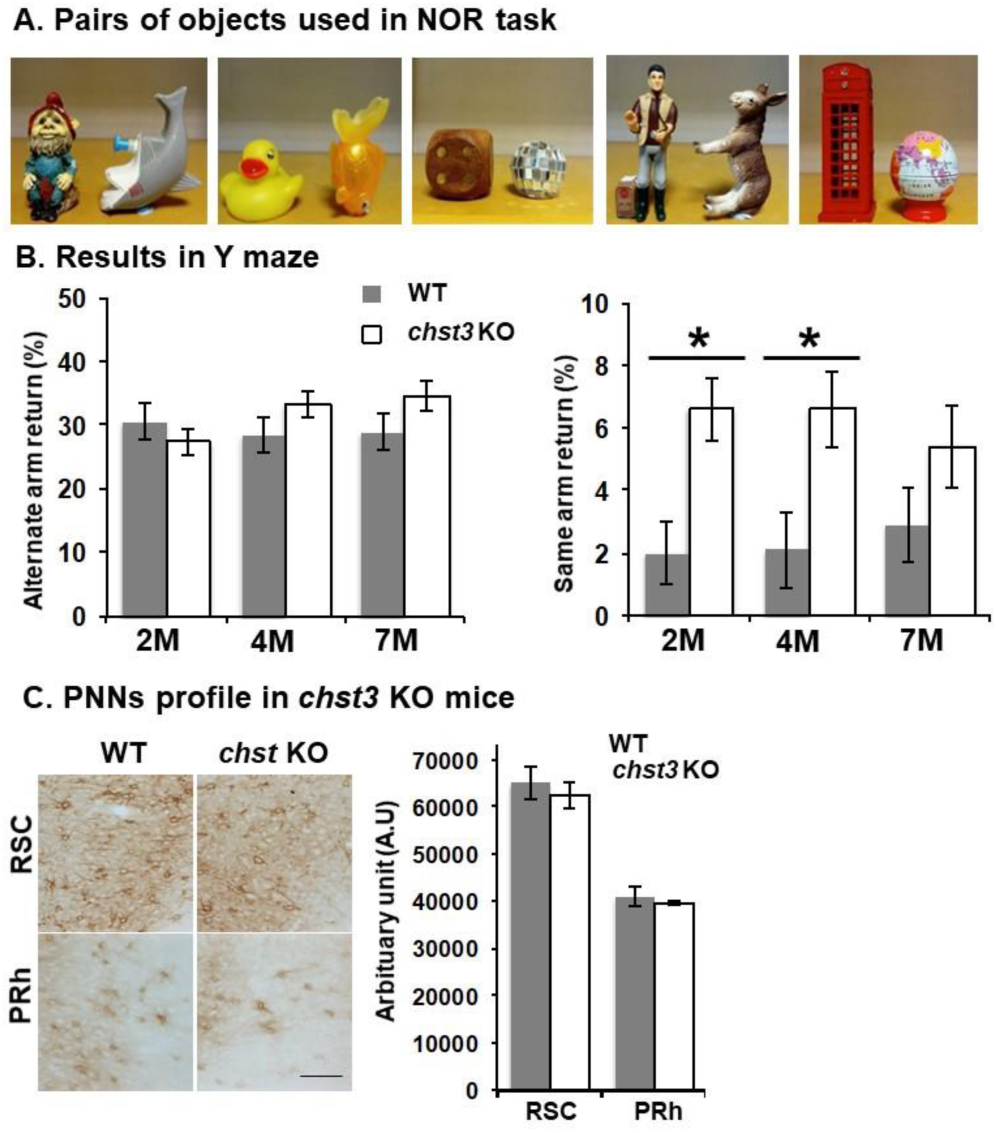
(A) Pairs of objects used in SOR task. (B) Y-maze based spontaneous alternation test in Chst3 KO mice at different ages, 2M: WT n=8, chst3 KO n=16, 4M: WT n=9, chst3 KO n=11, 7M: WT n=12, chst3 KO n=12. Left: Alternate arm return, Right: unpaired two-tailed t-test 2M: *p=0.018 4M: *p=0.019, 7M: n.s. p=0.2 (C) Analysis of PNNs in chst3 KO mice, Left: Immunohistochemistry of WFA(+) profile in retrosplenial cortex (RSC) and perirhinal cortex (PRh). Scale bar: 50 µm Right: WFA intensity comparison between WT and chst KO mice. There is no significant difference between WT and chst3 KO. n=6/group, Data represents as mean ± SEM,

**Supplementary Fig.3.**
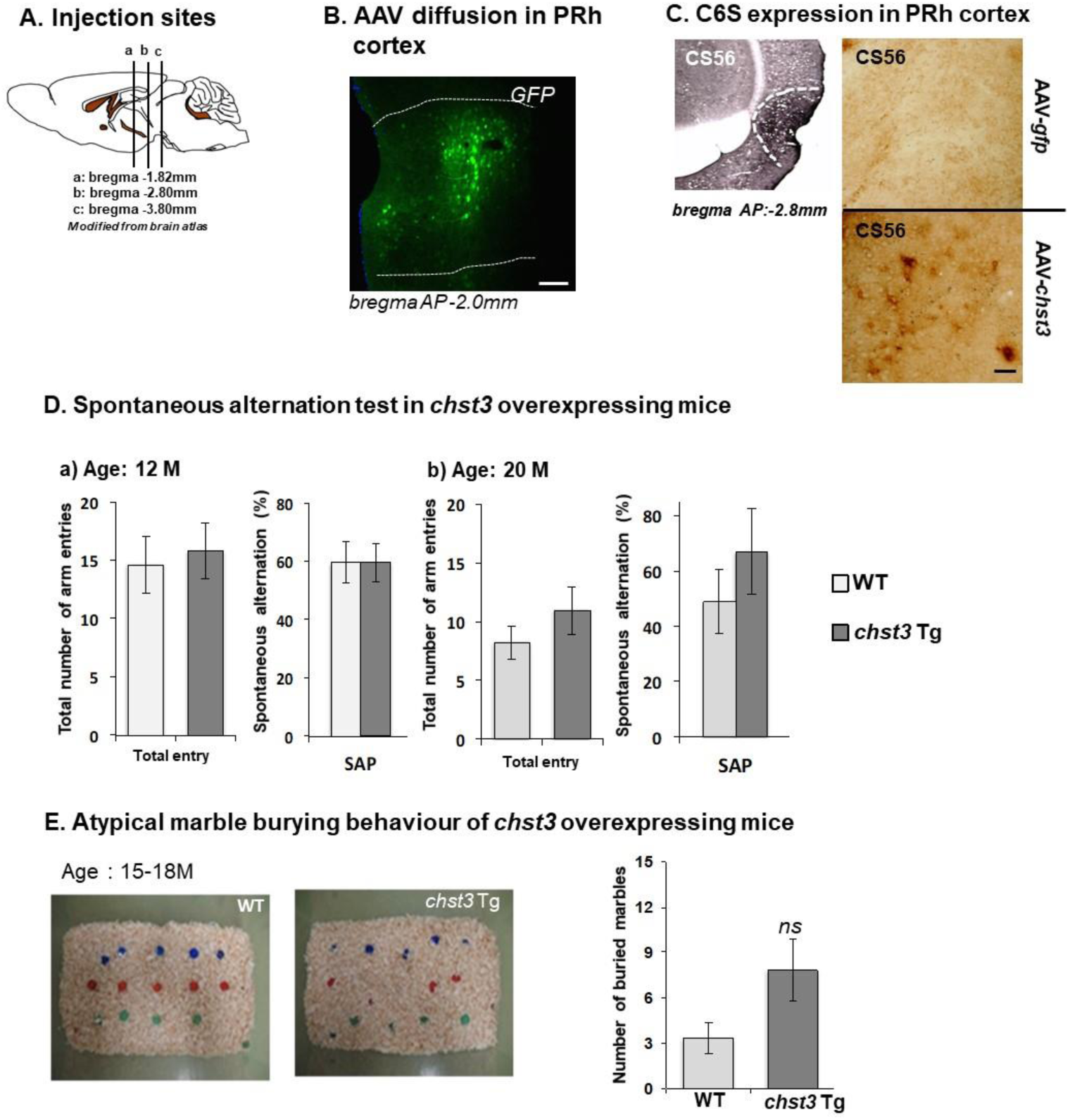
(A) Experimental time line of AAV-*chst3* injection to *chst3* KO mice at 4M. The viral vectors were injected to the perirhinal cortex and OR memory was tested at 4-5 weeks post injection. (B) The profile of AAV-GFP diffusion in perirhinal cortex, Scale bar: 100 µm (C) Experimental time line of AAV-*chst3* injection to 5M or 19M C57bl/6 mice. The viral vectors were injected to the perirhinal cortex and OR memory was tested at 4-5 weeks post injection. (D) C6S expression induced by AAV-*chst3* was detected by CS56 antibody in WT mice at 20M. AAV-GFP injection has no CS56 positive staining. Scale bar: 50 µm (E) Spontaneous alternation test in *chst3* Tg mice Age: 12M, WT n=5, *chst3* Tg n=9. Age: 20M, WT n=4 and *chst3* Tg n=4. There is no difference between WT and *chst3* Tg mice at 12M and 20M. (F) Marble burying behaviour of *chst3* Tg mice at the age of 15-18M. Left: representative images of marble burying test Right: WT n=6, *chst3* Tg n=6

